# The structure of probabilistic networks

**DOI:** 10.1101/016485

**Authors:** T. Poisot, A.R. Cirtwill, K. Cazelles, D. Gravel, M.-J. Fortin, D.B. Stouffer

## Abstract

1. There is a growing realization among community ecologists that interactions between species vary across space and time, and that this variation needs be quantified. Our current numerical framework to analyze the structure of species interactions, based on graph-theoretical approaches, usually do not consider the variability of interactions. Since this variability has been show to hold valuable ecological information, there is a need to adapt the current measures of network structure so that they can exploit it.
2. We present analytical expressions of key measures of network structured, adapted so that they account for the variability of ecological interactions. We do so by modeling each interaction as a Bernoulli event; using basic calculus allows expressing the expected value, and when mathematically tractable, its variance. When applied to non-probabilistic data, the measures we present give the same results as their non-probabilistic formulations, meaning that they can be generally applied.
3. We present three case studies that highlight how these measures can be used, in re-analyzing data that experimentally measured the variability of interactions, to alleviate the computational demands of permutationbased approaches, and to use the frequency at which interactions are observed over several locations to infer the structure of local networks. We provide a free and open-source implementation of these measures.
4. We discuss how both sampling and data representation of ecological networks can be adapted to allow the application of a fully probabilistic numerical network approach.

## Introduction

Ecological networks efficiently represent biotic interactions between individuals, populations, or species. Historically, their study focused on describing their structure, with a particular attention on food webs (Dunne 2006) and plant-pollinator interactions (Jordano 1987; Bascompte *et al.* 2003). This established that network structure is linked to community or ecosystem-level properties such as stability (McCann 2014), coexistence (Bastolla *et al.* 2009; Haerter *et al.* 2014), or ecosystem functioning (Duffy 2002). The description of ecological networks resulted in the emergence of questions about how functions and properties of communities emerged from their structure, and this stimulated the development of a wide array of measures for key network properties (Bersier *et al.* 2002; Banašek-Richter *et al.* 2004; Jordano & Bascompte 2013).

Given a network (*i.e.* a structure where nodes, most often species, are linked by edges, representing ecological interactions) as input, measures of network structure return a *property* based on one or several *units* (*e.g.* nodes, links, or groups thereof) from this network, either directly measured, or after an optimization process. Some of the properties are *direct* properties (they only require knowledge of the unit on which they are applied), whereas others are *emergent* (they require knowledge of, and describe, higher-order structures). For example, connectance, the realized proportion of potential interactions, is a direct property of a network, since it can be derived from the number of nodes and edges only. The degree of a node (how many interactions it is involved in) is a direct property of the node. The nestedness of a network (that is, the extent to which specialists and generalists overlap), is an emergent property, as it is not directly predictable from the degree of all nodes. The difference is no mere semantics: the difference between direct and emergent properties is important when interpreting their values. Direct properties are conceptually equivalent to means, in that they tend to be the first moment of network units, whereas emergent properties are conceptually equivalent to variances, higher-order moments, or probability distributions.

The interpretation of the measures of network structure as indicators of the action of ecological or evolutionary processes must now account for the numerous observations that network structure varies through space and time. In addition to the already well-established variation in the composition of the local species pool (Havens 2015), networks vary because species do not interact in a consistent way (Poisot *et al.* 2012).

Empirical evidence suggests that the network is not the right unit to understand this variation; rather, network variation emerges from the response of interactions to environmental factors and chance events (see Poisot *et al.* 2015 for a review). Interactions can vary for multiple (non-exclusive) reasons. Local mismatching in phenology creates *forbiden links* (Olesen *et al.* 2011; Vizentin-Bugoni *et al.* 2014; Maruyama *et al.* 2014). Local variations in abundance prevent the species from encountering one another (Canard *et al.* 2014). The joint action of neutral, phenologic, and behavioral effects, creates complex and hard to predict responses (Chamberlain *et al.* 2014; Olito & Fox 2015; Trøjelsgaard *et al.* 2015). For example, Olito & Fox (2015) showed that accounting for neutral (population-size driven) and trait-based effects allows the prediction of the cumulative change in network structure, but not of the change at the level of individual interactions. In addition, Carstensen *et al.* (2014) showed that not all interactions are equally variable within a meta-community: some are highly consistent, whereas others are extremely rare. These results suggest that species interactions, because they vary, cannot be adequately represented as yes-no events; it is therefore necessary to represent them as probabilities. We should replace the question of *Do these two species interact?* by *How likely is it that they will interact?*.

Yet the current way of dealing with probabilistic interactions are either to ignore variability entirely, or to generate networks with yes/no interactions based on the measured probabilities. Both approaches incur a net loss of information, and measures of network structure that explicitly account for interaction variability are a much needed mathematically rigorous alternative. When ignoring the probabilistic nature of interactions (henceforth *binary* networks), every non-zero element of the network is explicitly assumed to occur with probability 1. This over-represents rare events, and increases the number of interactions; as a result, this changes the estimated value of different network properties, in a way that remains poorly understood. The generation of random binary networks based on probabilities also suffers from biases, especially in the range of connectance within which most ecological systems lie. These biases are (i) pseudo-replication when the permutational space is small (Poisot & Gravel 2014), and (ii) systematic biases in the emergent properties at low connectances (Chagnon 2015). An alternative is to consider only the interactions above a given threshold, which unfortunately leads to under-representation of rare events and decreases the effective number of interactions. The use of thresholds also notably introduces the risk of removing species that have a lot of interactions that individually have a low probability of occurring. These considerations highlight the need to amend our current methodology for the description of ecological networks, in order to give more importance to the variation of individual interactions.

Yet the extant methodological corpus is well accepted, and the properties it describes are well understood. Rather than suggesting measures, we argue that it is more productive to re-express those we already have, in a way that does not lose information when applied to probabilistic networks. We contribute to this effort by re-developing a unified toolkit of measures to characterize the structure of probabilistic interaction networks. Several direct and emergent core properties of ecological networks (both bipartite and unipartite) can be re-formulated in a probabilistic context. We illustrate this toolkit through several case studies, and discuss how the current challenges in the (i) measurement and (ii) analysis of probabilistic interaction networks.

## Suite of probabilistic network metrics

We use the following notation throughout the paper. **A** is a matrix where each element *A*_*ij*_ gives P(*ij*), *i.e.* the probability that species *i* establishes an interaction with species *j*. If **A** represents a unipartite network (*e.g.* a food web), it is a square matrix and contains the probabilities of each species interacting with all others, including itself. If **A** represents a bipartite network (*e.g.* a pollination network), it will not necessarily be square. We call *S* the number of species, and *R* and *C* respectively the number of rows and columns. *S* = *R* = *C* in unipartite networks, and *S* = *R* + *C* in bipartite networks. Note that all of the measures defined below can be applied on a bipartite network that has been made unipartite. The unipartite transformation of a bipartite matrix **A** is the block matrix **B**:

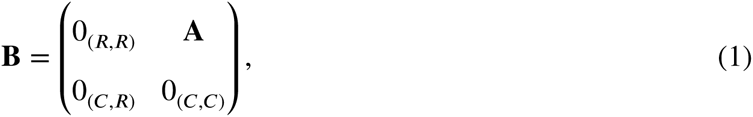

where 0_(*C,R*)_ is a matrix of *C* rows and *R* columns (noted *C* × *R*) filled with 0s, etc. Note that for centrality to be relevant in bipartite networks, this matrix should be made symmetric: **B**_*ij*_ = **B**_*ji*_.

We assume that all interactions are independent (so that P(*ij* ∩ *kl*) = P(*ij*)P(*kl*) for any species), and can be represented as a series of Bernoulli trials (so that 0 ≤ P(*ij*) ≤ 1). A Bernoulli trial is the realization of a probabilistic event that gives 1 with probability P(*ij*) and 0 otherwise. The latter condition allows us to derive estimates for both the *variance* (var(*X*) = *p*(1 - *p*)) and expected values (E(*X*) = *p*) of the network measures. The variance of additive independent events is the sum of their individual variances, and the variance of multiplicative independent events is

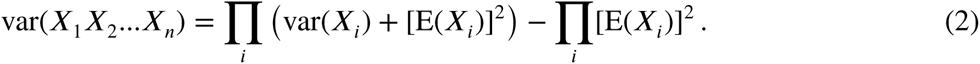

As all *X*_*i*_ are Bernoulli random variables,

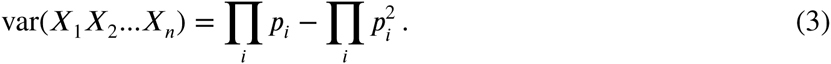

As a final note, all of the measures described below can be applied on the binary (0/1) versions of the networks in which case they converge on the non-probabilistic version of the measure as usually calculated. This property is particularly desirable as it allows our framework to be used on any unweighted network represented in a probabilistic or binary way. The approach outlined here differs from using *weighted* networks, in that it answers a different ecological question. Probabilistic networks describe the probability that any interaction will happen, whereas weighted networks describe some measure of the effect of the interaction when it happens (Berlow *et al.* 2009); weighted networks therefore assume that the interaction happen. Although there are several measures for weighted ecological networks (Bersier *et al.* 2002), in which interactions happen but with different outcomes, these are not relevant for probabilistic networks; they do not account for the fact that interactions display a variance that will cascade up to the network level. Instead, the weight of each interaction is best viewed as a second modeling step focusing on the non-zero cases (*i.e.* the interactions that are realized); this is similar to the method now frequently used in species distribution models, where the species presence is modeled first, and its abundance second, using a (possibly) different set of ecological predictors (Boulangeat *et al.* 2012).

## Direct network properties

### Connectance and number of interactions

Connectance (or network density) is the proportion of possible interactions that are realized, defined as *Co* = *L*/(*R*×*C*), where *L* is the total number of interactions. As all interactions in a probabilistic network are assumed to be independent, the expected value of *L*, is

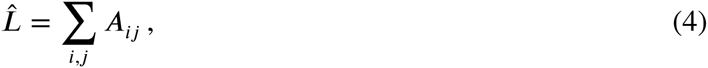

and 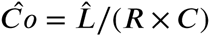. Likewise, the variance of the number of interactions is 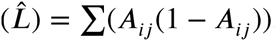.

### Node degree

The degree distribution of a network is the distribution of the number of interactions established (number of successors) and received (number of predecessors) by each node. The expected degree of species *i* is

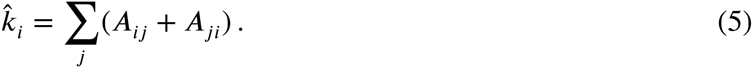

The variance of the degree of each species is 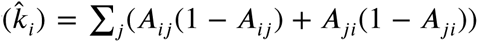. Note also that 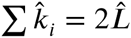, as expected

### Generality and vulnerability

By simplification of the above, generality *g*_*i*_ and vulnerability *v*_*i*_ are given by, respectively,∑ *j A*_*ij*_ and ∑ *j A*_*ji*_, with their variances ∑ *j A*_*ij*_(1 - *A*_*ij*_) and ∑ *j A*_*ji*_(1 - *A*_*ji*_).

## Emergent network properties

### Path length

Networks can be used to describe indirect interactions between species through the use of paths. The existence of a path of length 2 between species *i* and *j* means that they are connected through at least one additional species *k*. In a probabilistic network, unless some elements are 0, all pairs of species *i* and *j* are connected through a path of length 1, with probability *A*_*ij*_. The expected number of paths of length *k* between species *i* and *j* is given by

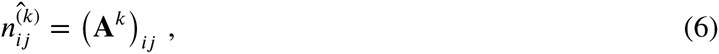

where **A**^*k*^ is the matrix multiplied by itself *k* times.

It is possible to calculate the probability of having at least one path of length *k* between the two species: this can be done by calculating the probability of having no path of length *k*, then taking the running product of the resulting array of probabilities. For the example of length 2, species *i* and *j* are connected through *g* with probability *A*_*ig*_*A*_*gj*_, and so this path does not exist with probability 1 - *A*_*ig*_*A*_*gj*_. For any pair *i*, *j*, let **m** be the vector such that *m*_*g*_ = *A*_*ig*_*A*_*gj*_ for all *g* ∉ (*i, j*) (Mirchandani 1976). The probability of not having any path of length 2 is (Π1 - **m**). Therefore, the probability of having a path of length 2 between *i* and *j* is

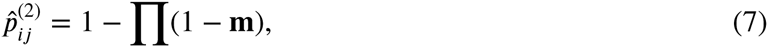

which can also be noted

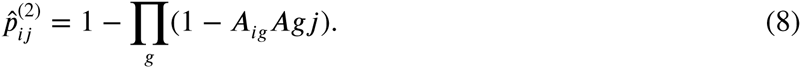

In most situations, one would be interested in knowing the probability of having a path of length 2 *without* having a path of length 1; this is simply expressed as 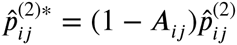. These results can be expanded to any length *k* in [2, *n* - 1]. First one can, by the same logic, generate the expression for having at least one path of length *k*:

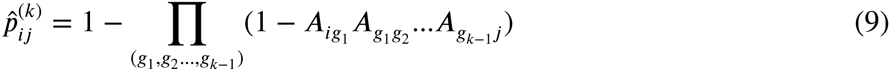

where (*g*_1_, *g*_2_*…, g*_*k*-1_) are all the (*k* - 1)-permutations of 1, 2, *…, n*\(*i, j*). Then having a path of length *k* without having any smaller path is

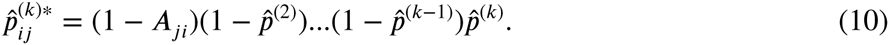

### Unipartite projection of bipartite networks

The unipartite projection of a bipartite network is obtained by linking any two nodes of one mode (“side” of the network) that are connected through at least one node of the other mode; for example, two plants are connected if they share at least one pollinator. It is readily obtained using the formula in the *Path length* section. This yields either the probability of an edge in the unipartite projection (of the upper or lower nodes), or if using the matrix multiplication, the expected number of such nodes.

### Nestedness

Nestedness is an important measure of (bipartite) network structure that tells the extent to which the interactions of specialists and generalists overlap. We use the formula for nestedness proposed by Bastolla *et al.* (2009); this measure is a modification of NODF (Almeida-Neto *et al.* 2008) for ties in species degree that removes the constraint of decreasing fill. Nestedness for each margin of the matrix is defined as *χ*^(*R*)^ and *η*^(*C*)^ for, respectively, rows and columns. As per Almeida-Neto *et al.* (2008), we define a global statistic for nestedness as *η*= (*η*^(*R*)^ + *η*^(*C*)^)/2.

Nestedness, in a probabilistic network, is defined as

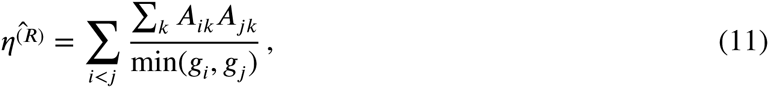

where *g*_*i*_ is the expected generality of species *i*. The reciprocal holds for *η*^(*C*)^ when using *v*_*i*_ (the vulnerability) instead of *g*_*i*_. The values returned are within [0; 1], with η = 1 indicating complete nestedness.

### Modularity

Modularity represents the extent to which networks are compartmentalized, *i.e.* the tendency for subsets of species to be strongly connected together, while they are weakly connected to the rest of the network (Stouffer & Bascompte 2011). Modularity is measured as the proportion of interactions between nodes of an arbitrary number of modules, as opposed to the random expectation.The modularity as derived by Newman (2004) can be expressed as

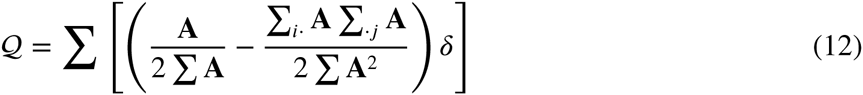

where ∑_*i*·_ **A** and ∑_·*j*_ **A** are the sums of rows and columns of **A**, and *δ* is a matrix, wherein *δ*_*ij*_ is 1 if *i* and *j* belong to the same module, and 0 otherwise. This formula can be *directly* applied to probabilistic networks. Modularity takes values in [0; 1], where 1 indicates perfect modularity.

### Centrality

Although node degree is a rough first order estimate of centrality, other measures are often needed. Here, we derive the expected value of centrality according to Katz (1953). This measure generalizes to directed acyclic graphs (whereas other do not). For example, although eigenvector centrality is often used in ecology, it cannot be measured on probabilistic graphs. Eigenvector centrality requires the matrix’s largest eigenvalues to be real, which is not the case for all probabilistic matrices. The measure proposed by Katz is a useful replacement, because it accounts for the paths of all length between two species instead of focusing on the shortest path.

As described above, the expected number of paths of length *k* between *i* and *j* is (**A**^*k*^)_*ij*_. Based on this, the expected centrality of species *i* is

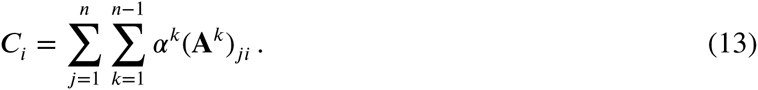

The parameter *α* ∈ [0; 1] regulates how important long paths are. When *α* = 0, only first-order paths are accounted for (and the centrality is equal to the degree). When *α* = 1, paths of all length are equally important. As *C*_*i*_ is sensitive to the size of the matrix, we suggest normalizing by **C** = ∑*C* so that

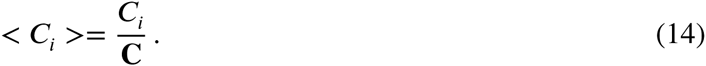

This results in the *expected relative centrality* of each node in the probabilistic network, which sums to unity.

### Species with no outgoing links

Estimating the number of species with no outgoing links (successors) can be useful when predicting whether, *e.g.*, predators will go extinct. Alternatively, when prior information about traits are available, this can allows predicting the invasion success of a species in a novel community.

A species has no successors if it manages *not* to establish any outgoing interaction, which for species *i* happens with probability

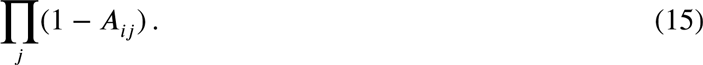

The number of expected such species is therefore the sum of the above across all species,

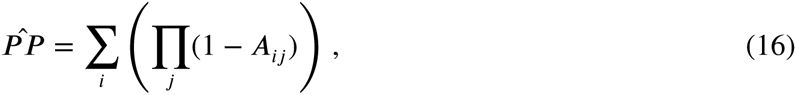

and its variance is

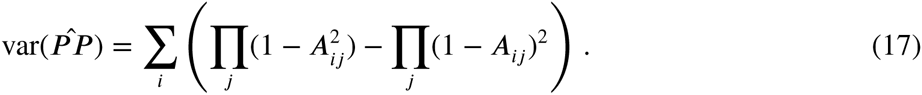

Note that in a non-probabilistic context, species with no outgoing links would be considered primary producers. This is not the case here: if interactions are probabilistic events, then even a top predator may have no preys, and this clearly doesn’t imply that it will become a primary producer in the community. For this reason, the trophic position of the species may be measured better with the binary version of the matrix.

### Species with no incoming links

Using the same approach as for the number of species with no outgoing links, the expected number of species with no incoming links is therefore

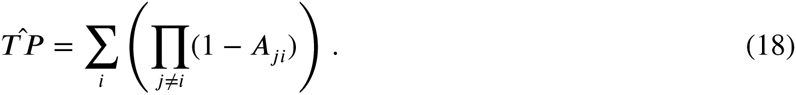

Note that we exclude self-interactions, as top-predators in food webs can, and often do, engage in cannibalism.

### Number of species with no interactions

Predicting the number of species with no interactions (or whether any species will have at least one interaction) is useful when predicting whether species will be able to integrate into an existing network, for example. From a methodological point of view, this can also be a helpful *a priori* measure to determine whether null models of networks will have a lot of species with no interactions, and so will require intensive sampling.

A species has no interactions with probability

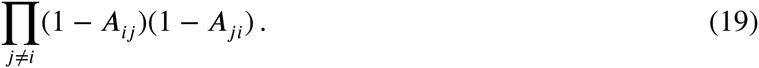

As for the above, the expected number of species with no interactions (*free species*) is the sum of this quantity across all *i*:

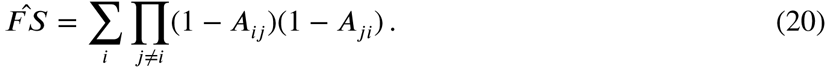

The variance of the number of species with no interactions is

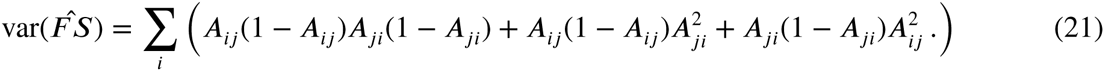

### Self-loops

Self-loops (the existence of an interaction of a species onto itself) is only meaningful in unipartite networks. The expected proportion of species with self-loops is very simply defined as Tr(**A**), that is, the sum of all diagonal elements. The variance is Tr(**A** ◊ (1 - **A**)), where ◊ is the element-wise product operation (Hadamard product).

### Motifs

Motifs are sets of pre-determined interactions between a fixed number of species (Milo *et al.* 2002; Stouffer *et al.* 2007), such as apparent competition with one predator sharing two prey. As there are an arbitrarily large number of motifs, we will illustrate the approach with only two examples.

The probability that three species form an apparent competition motif where *i* is the predator, *j* and *k* are the prey, is

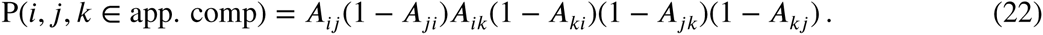

Similarly, the probability that these three species form an omnivory motif, in which *i* and *j* consume *k* and *i* consumes *j*, is

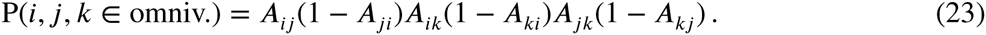

The probability of the number of *any* three-species motif motif m in a network is given by

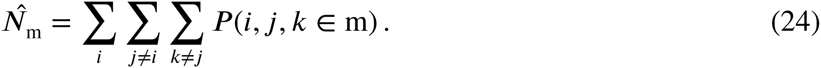

It is indeed possible to have an expression of the variance of this value, or of the variance of any three species forming a given motif, but their expressions become rapidly untractable and are better computed than written.

## Network comparison

The dissimilarity of a pair of (ecological) networks can be measured using the framework set forth by Koleff *et al.* (2003) using *β*-diversity measures. Measures of *β*-diversity compute the dissimilarity between two networks based on the cardinality of three sets, *a*, *c*, and *b*, which are respectively the shared items, items unique to superset (network) 1, and items unique to superset 2 (the identity of which network is 1 or 2 matters for asymmetric measures). Supersets can be the species within each network, or the interactions between species. Following Poisot *et al.* (2012), the dissimilarity of two networks can be measured as either *β*_*W* *N*_ (all interactions), or *β*_*OS*_ (interactions involving only common species), with *β*_*OS*_ ≤ *β*_*OS*_.

Within our framework, these measures can be applied to probabilistic networks. The expected values of *ā*, *c̄*, and *b̄*are, respectively, ∑**A**_1_ ◊ **A**_2_, ∑**A**_1_ ◊ (1 - **A**_2_), and ∑(1 - **A**_1_) ◊ **A**_2_. Whether *β*_*OS*_ or *β*_*W* *N*_ is measured requires to alter the matrices **A**_1_ and **A**_2_. To measure *β*_*OS*_, one must remove all unique species; to measure *β*_*W* *N*_, one must expand the two matrices so that they have the same species at the same place, and give a weight of 0 to the added interactions.

## Implementation

We provide these measures of probabilistic network structure in a free and open-source (MIT licensed) library for the julia language, available at http://github.com/PoisotLab/EcologicalNetwork.jl. The code can be cited using the following DOI: 10.5281/zenodo.28317 (version 1.0.1). A user guide, including examples, resides at http://ecologicalnetworkjl.readthedocs.org/.

### Case studies

In this section, we contrast the use of probabilistic measures to the current approaches of either using binary networks, or working with null models through simulations. When generating random networks, what we call *Bernoulli trials* from here on, a binary network is generated by doing a Bernoulli trial with probability *A*_*ij*_, for each element of the matrix. This generates networks that have only 0/1 interactions, and are realizations of the probabilistic network. This is problematic because higher order structures involving rare events will be under-represented in the sample, and because most naive approaches (*i.e.* not controlling for species degree) are likely to generate species with no interactions, especially in sparsely connected networks frequently encountered in ecology (Milo *et al.* 2003; Poisot & Gravel 2014; Chagnon 2015) – on the other hand, non-naive approaches (*e.g.* based on swaps or quasi-swaps) break the assumption of independence between interactions.

### Comparison of probabilistic networks

In this sub-section, we apply the above probabilistic measures to a bacteria–phage interaction network. Poullain *et al.* (2008) measured the probability that 24 phage can infect 24 strains of bacteria of the *Pseudomonas fluorescens* species (group SBW25). The (probabilistic) adjacency matrix was constructed by estimating the probability of each phage–bacteria interaction though independent infection assays, and can take values of 0, 0.5 (interaction is variable), and 1.0. We have generated a “Binary” network by setting all interactions with a probability higher than 0 to unity, to simulate the results that would have been obtained in the absence of estimates of interaction probability.

Measuring the structure of the Binary, Bernoulli trials, and Probabilistic network gives the following results (average, and variance when there is an analytical expression):

| Measure | Binary | Bernoulli trials | Probabilistic |
| --- | --- | --- | --- |
| links | 336 | $221.58 \pm 57.57$ | $221.52 \pm 57.25$ |
| $\eta$ | 0.73 | 0.528 | 0.512 |
| $\eta^{(R)}$ | 0.72 | 0.525 | 0.507 |
| $\eta^{(C)}$ | 0.75 | 0.531 | 0.518 |
| one consumer, two resources motif | 4784 | 2089 | 2110 |
| two consumers, one resource motif | 4718 | 2116 | 2120 |

As these results show, treating all interactions as having the same probability, *i.e.* removing the information about variability, (i) overestimates nestedness by ≈ 0.2, (ii) overestimates the number of links by 115, and (iii) overestimates the number of motifs (we have limited our analysis to the two following motifs: one consumer sharing two resources, and two consumers competing for one resource). For the number of links, both the probabilistic measures and the average and variance of 10^4^ Bernoulli trials were in strong agreement (they differ only by the second decimal place). For the number of motifs, the difference was larger, but not overly so. It should be noted that, especially for computationally demanding operations such as motif counting, the difference in runtime between the probabilistic and Bernoulli trials approaches can be extremely important.

Using Bernoulli trials had the effect of slightly over-estimating nestedness. The overestimation is statistically significant from a purely frequentist point of view, but significance testing is rather meaningless when the number of replicates is this large and can be increased arbitrarily; what is important is that the relative value of the error is small enough that Bernoulli trials are able to adequately reproduce the probabilistic structure of the network. It is not unexpected that Bernoulli trials are this close to the analytical expression of the measures; due to the experimental design of the Poullain *et al.* (2008) study, probabilities of interactions are bound to be high, and so variance is minimal (most elements of **A** have a value of either 0 or 1, and so their individual variance is 0 – though their confidence interval varies as a function of the number of observations from which the probability is derived). Still, despite overall low variance, the binary approach severely mis-represents the structure of the network.

### Null-model based hypothesis testing

In this section, we analyse 59 pollination networks from the literature using two usual null models of network structure, and two models with intermediate constraints. These data cover a wide range a situations, from small to large, and from densely to sparsely connected networks. They provide a good demonstration of the performance of probabilistic metrics. Data come from the *InteractionWeb Database*, and were queried on Nov. 2014.

We use the following null models. First (Type I, Fortuna & Bascompte (2006)), any interaction between plant and animals happens with the fixed probability P = *Co*. This model controls for connectance, but removes the effect of degree distribution. Second, (Type II, Bascompte *et al.* (2003)), the probability of an interaction between animal *i* and plant *j* is (*k*_*i*_/*R* + *k*_*j*_/*C*)/2, the average of the richness-standardized degree of both species. In addition, we use the models called Type III in and out (Poisot *et al.* 2013), that use the row-wise and column-wise probability of an interaction respectively, as a way to understand the impact of the degree distribution of upper and lower level species.

Note that these null models will take a binary network and, through some rules turn it into a probabilistic one. Typically, this probabilistic network is used as a template to generate Bernoulli trials and measure some of their properties, the distribution of which is compared to the empirical network. This approach is computationally inefficient (Poisot & Gravel 2014), especially using naive models (Milo *et al.* 2003), and as we show in the previous section, can yield biased estimates of the true average of nestedness (and presumably other properties).

We measured the nestedness of the 59 (binary) networks, then generated the random networks under the four null models, and calculated the expected nestedness using the probabilistic measure. Our results are presented in Figure 1.

**Figure 1:**
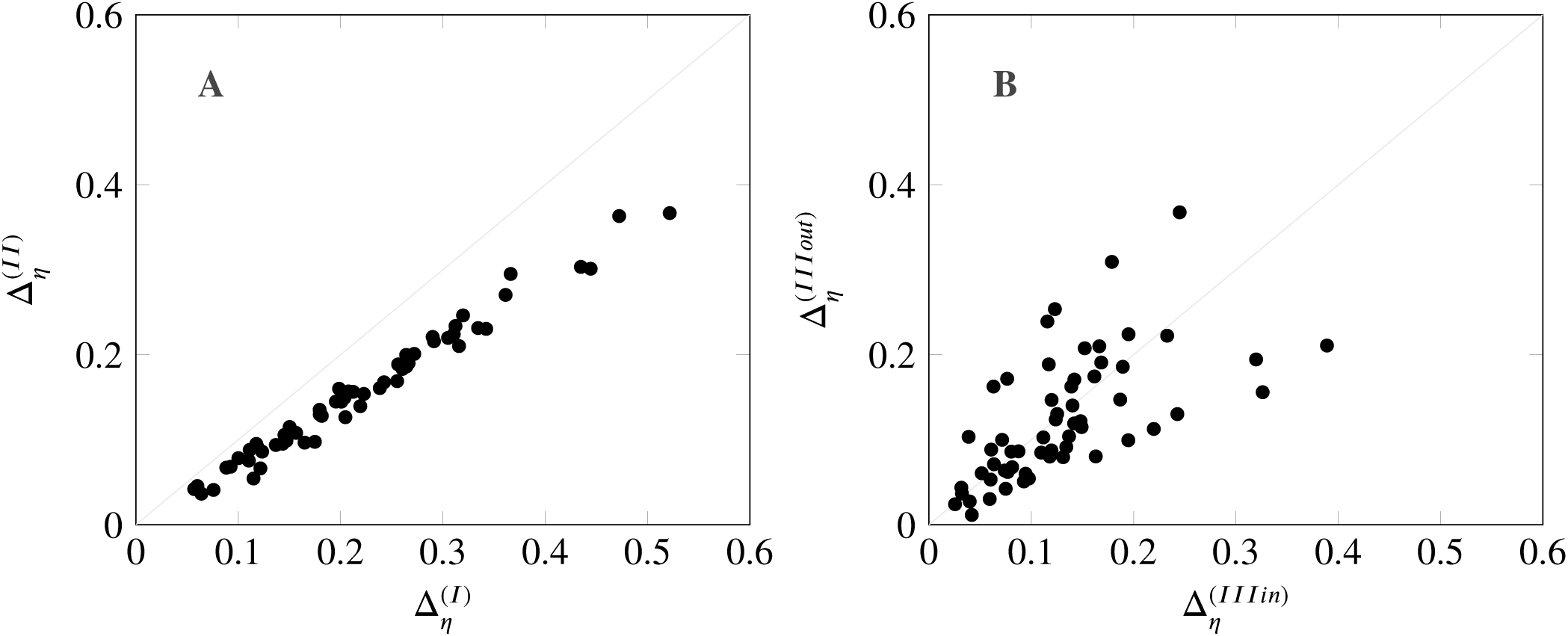
Results of the null model analysis of 59 plant-pollination networks. **A**. There is a consistent tendency for (i) both models I and II to estimate less nestedness than in the empirical network, although null model II yields more accurate estimates. **B**. Models III in and III out also estimate less nestedness than the empirical network, but neither has a systematic bias. For each null model *i*, the difference 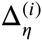 in nestednessχis expressed as 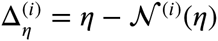, where.N ^(*i*)^(*1*) is the nestedness of null model *i*.

There are two striking results. First, empirical data are consistently *more* nested than the null expectation, as evidenced by the fact that all Δ_*ij*_ values are strictly positive. Second, this underestimation is *linear* between null models I and II, although null model II is always closer to the nestedness of the empirical network (which makes sense, since null model II incorporates the higher order constraint of approximating the degree distribution of both levels). That the nestedness of the null model probability matrix is so strongly determined by the nestedness of the empirical networks calls for a closer evaluation of how the results of null models are interpreted (especially since networks generated using Bernoulli trials revealed a very low variance in their nestedness).

There is a strong, and previously unaccounted for, circularity in this approach: empirical networks are compared to a null model which, as we show, has a systematic bias *and* a low variance (in the properties of the networks it generates), meaning that differences in nestedness that are small (thus potentially ecologically irrelevant) have a good chance of being reported as significant. Interestingly, models III in and III out made overall *fewer* mistakes at estimating nestedness – respectively 0.129 and 0.123, compared to resp. 0.219 and 0.156 for model I and II. Although the error is overall sensitive to model type (KruskalWallis χ^2^ = 35.80, d.f. = 3, *p* 10^-4^), the three pairs of models that where significantly different after controlling for multiple comparisons are I and II, I and III in, and I and III out (model II is not different from either models III in or out).

In short, this analysis reveals that (i) the null expectation of a network property under randomization scenarios can be obtained through the analysis of the probabilistic matrix, instead of the analysis of simulated Bernoulli networks; (ii) different models have different systematic biases, with models of the type III performing better for nestedness than any other models. This can be explained by the fact that nestedness of a network, as expressed by Bastolla *et al.* (2009), is the average of a row-wise and column-wise nestedness. These depend on the species degree, and as such should be well predicted by models III. The true novelty of the approach outlined here is that, rather than having to calculate the measure for thousands of replicates, an *unbiased* estimate of its mean can be obtained in a fraction of the time using the measures described here. This is particularly important since, as demonstrated by Chagnon (2015), the generation of null randomization is subject to biases in the range of connectance where most ecological networks fall. Our approach aims to provide a bias-free, time-effective way of estimating the expected value of a network property.

### Spatial-variation predicts local network structure

In this final application, we re-analyze data from a previous study by Trøjelsgaard *et al.* (2015), to investigate how spatial information can be used to derive probability of interactions. In the original dataset, fourteen locations have been sampled to describe the local plant-pollination network. This dataset exhibits both species and interaction variability across sampling locations. We define the overall probability of an interaction in the following way,

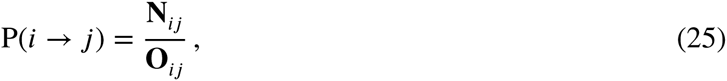

where **O**_*ij*_ is the number of sampling locations in which both pollinator *i* and plant *j* co-occur, and **N**_*ij*_ is the number of sampling locations in which they interact. This takes values between 0 (no co-occurence *or* no interactions) and 1 (interaction observed every time there is co-occurrence, including single observations of an interacting species pair). This represents a simple probabilistic model, in which it is assumed that our ability to observe the interaction is a proxy of how frequent it is.

Based on this information, we compare the connectance, nestedness, and modularity, of each sampled (binary) network, to the expected values if interactions are well predicted by the probability given above. The results are presented in Figure 2. There is a clear linear, positive correlation (coeff. 0.89 for connectance, 0.76 for *η*, and 0.92 for modularity) between the observed network properties (binary matrices) and the predictions based on the probabilistic model. This analysis, although simple, suggest that the *local* structure of ecological networks can represent the outcome of a filtering of species interactions, the signature of which can be detected at the regional level by a variation in the probabilities of interactions. Note however that this approach *does not* allow predicting the structure of any arbitrary species pool, since it cannot know the probability of an interaction between two species that never co-occured.

**Figure 2:**
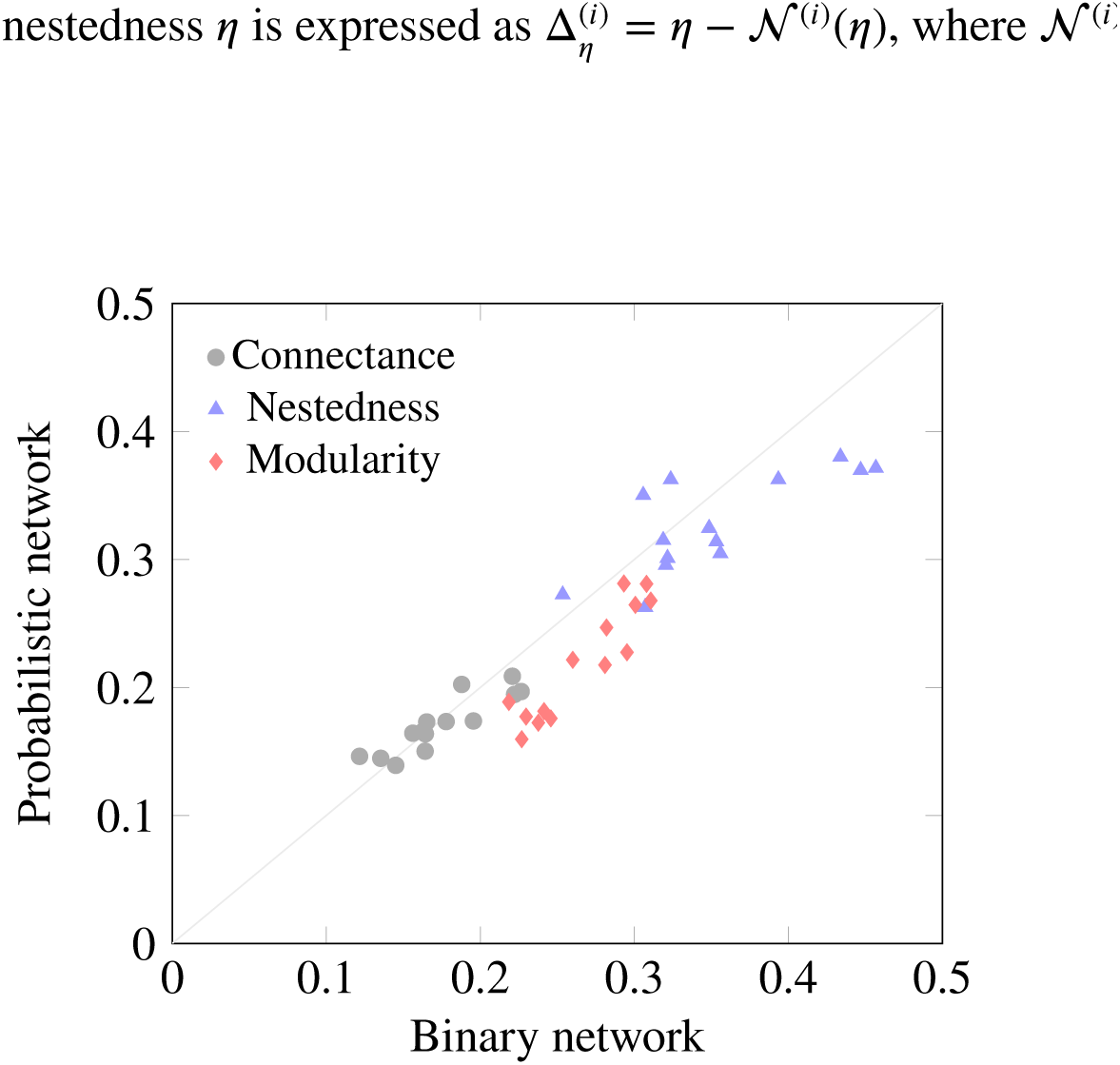
Local network structure infered from the locally observed interactions (x-axis) or the spatial probabilistic model (y-axis) in the Canaria Island dataset. Although the binary networks slightly underestimate the properties studied here, there is a positive and linear relationship between the empirical structure, and the structure predicted based on probabilities of interactions derived from occurrence information. tools that address all forms of complexity, the most oft-neglected yet pervasive of which is the fact that interactions are variable. Through the suite of measures we present here, we allow future analyses of network structure to account for this phenomenon. There are two main considerations highlighted by this methodological development. First, in what way are probabilistic data are actually independent? Second, what are the implications for data collection?

## Discussion

Understanding the structure of ecological networks, and whether it relates to emergent ecosystem properties, is a strong research agenda for community ecology. A proper estimation of this structure requires tools that address all forms of complexity, the most 1 oft-neglected yet pervasive of which is the fact that interactions are variable. Through the suite of measures we present here, we allow future analyses of network structure to account for this phenomenon. There are two main considerations highlighted by this methodological development. First, in what way are probabilistic data are actually independent? Second, what are the implications for data collection?

### Non-independance of interactions

We developed and presented a set of measures to quantify the expected network structure, using the probability that each interaction is observed or happens, in a way that does not require time-consuming simulations. Our framework is set up in such a way that the probabilities of interactions are considered to be independent. This is an over-simplification of what we understand of ecological reality, where interactions have effects on one another (Golubski & Abrams 2011; Sanders & Veen 2012; Ims *et al.* 2013). Yet we feel that, as a first approximation, this assumption is reasonable. There is a strong methodological argument for which the non-independance of interactions cannot currently be robustly accounted for: analytical expectations for non-independant Bernoulli events require knowledge of the full dependence structure. Not only does this severely limit the ability to provide measures of network structure, it requires a far more extensive sampling that what is needed to obtain an estimate of the probability of interactions one by one.

### Estimates of interaction probabilities

Estimating interaction probabilities based on species abundances (Canard *et al.* 2014; Olito & Fox 2015) do not yield independent probabilities: changing the abundance of one species changes all probabilities in the network. They are not Bernoulli events either, as the sum of all probabilities derived this way sums to unity. On the other hand, “cafeteria experiments” (in which individuals from two species are directly exposed to one another to observe whether or not an interaction occurs) give truly independent probabilities of interactions—although this approach is limited to systems with a small number of species, and that are amenable to microcosms or mesocosms experiments. Using the approach outlined by Poisot *et al.* (2015), different sources of information (species abundance, trait distribution, and the outcome of experiments) can be combined to estimate the probability that interactions will happen in empirical communities.

Another way to obtain approximation of the probability of interactions is to use spatially and temporally replicated sampling (assuming that replicates are done in environments that can be assumed to be comparably homogeneous); in this context, it is not the interactions that are repeatedly sampled, but the network as a whole. Some studies (Tylianakis *et al.* 2007; Carstensen *et al.* 2014; Olito & Fox 2015; Trøjelsgaard *et al.* 2015) surveyed the existence of interactions at different locations, and a simple approach of dividing the number of observations of an interaction by the number of co-occurence of the species involved will provide a (somewhat crude) estimate of the probability of this interaction. This approach requires extensive sampling, especially since interactions are harder to observe than species (Poisot *et al.* 2012; Gilarranz *et al.* 2015), yet it enables the re-analysis of existing datasets in a probabilistic context.

### Implications for data collection

An important outcome is that, when estimating probabilities from observational data, it becomes possible to have an estimate of how robust the sampling is. How completely a network is sampled is a key, yet often-overlooked, driver of some measures of structure (Nielsen & Bascompte 2007; Chacoff *et al.* 2012; Fründ *et al.* 2015). The probabilistic approach allows to estimate the *confidence interval* of the interaction probability, knowing the number of samples used for the estimation. Assuming normally distributed observational error (this can be generalized for other error distributions), the confidence interval around a probability *p* estimated from *n* samples is

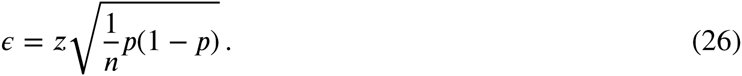

For a 95% confidence interval, *z ≈*1.96. If an interaction is estimated to happen at *p* = 0.3, its 95% confidence interval is [0; 0.74] when estimated from four samples, [0.01; 0.58] when estimated from ten, and [0.21; 0.38] when estimated from a hundred. Note that the above formula tends to perform poorly when *n <* 30, and do not applies when *p ϵ* {0, 1}; it nevertheless provides an *estimate* of how robust the probability estimate is.

The quantification, and integration, of uncertainty in the probability of interaction, is a subject that remains to be worked out. To develop a coarse understanding of how it affects the estimate of network properties, one can (for example) sample the interaction probability within its 95% confidence interval. This points to a fundamental issue with the sampling of networks: a precise estimate of the probability of interactions from observational data is tremendously difficult to achieve. Although the development of predictive models partly alleviates this difficulty, estimating confidence intervals around the probability of an interaction guide empirical research efforts to (i) either collect additional replicates or (ii) provide additional data to improve the performance of predictive models.

## Acknowledgements

This work was funded by a CIEE working group grant to TP, DG, and DBS. TP is funded by a starting grant from the Université de Montréal, and a Discovery Grant from NSERC. DBS acknowledges support from a Marsden Fund Fast-Start grant (UOC-1101) and Rutherford Discovery Fellowship, both administered by the Royal Society of New Zealand. The idea of network measures as direct/emergent properties of network units was first discussed during the *Web of Life* meeting, held in Montpellier in 2012.

